# Intercellular crosstalk in adult dental pulp is mediated by heparin-binding growth factors Pleiotrophin and Midkine

**DOI:** 10.1101/2022.10.28.514178

**Authors:** Natnicha Jiravejchakul, Gabriela L. Abe, Martin Loza, Soyoung Park, Ponpan Matangkasombut, Jun-Ichi Sasaki, Satoshi Imazato, Diego Diez, Daron M. Standley

**Affiliations:** Department of Genome Informatics, Research Institute for Microbial Diseases, Osaka University, 3-1 Yamadaoka, Suita 565-0871, Japan; Department of Microbiology, Faculty of Science, Mahidol University, Bangkok 10400, Thailand; Department of Advanced Functional Materials Science, Osaka University Graduate School of Dentistry, 1-8 Yamadaoka, Suita 565-0871, Japan; Laboratory of Functional Analysis in silico, Human Genome Center, The Institute of Medical Science, The University of Tokyo, 4-6-1 Shirokane-dai, Minato-ku, Tokyo 108-8639, Japan; Department of Systems Immunology, Immunology Frontier Research Institute, Osaka University, 3-1 Yamadaoka, Suita 565-0871, Japan; Department of Biomaterials Science, Osaka University Graduate School of Dentistry, 1-8 Yamadaoka, Suita 565-0871, Japan; Quantitative Immunology Research Unit, Immunology Frontier Research Institute, Osaka University, 3-1 Yamadaoka, Suita 565-0871, Japan

## Abstract

In-depth knowledge of the cellular and molecular composition of dental pulp (DP) and the crosstalk between DP cells that drive tissue homeostasis or regeneration are not well understood. To address these questions, we performed data analysis of publicly available single-cell transcriptomes of DP. This analysis revealed that DP resident fibroblasts have a unique gene expression profile when compared with fibroblasts from 5 other reference tissues: blood, bone marrow, adipose tissue, lung, and skin. Genes coded for heparin-binding growth-factors, pleiotrophin (PTN) and midkine (MDK), possessed the highest differential expression levels in DP fibroblasts. In addition, we identified extensive crosstalk between DP fibroblasts and several other DP cells, including Schwann cells, MSCs and odontoblasts. These findings suggest that fibroblast-derived growth factors regulate DP niches, and thus have a potential role as dental therapeutic targets.

## Introduction

Dental pulp (DP) is a soft tissue of ectomesenchymal origin, located within teeth, and surrounded by rigid dentine walls. Vascularization and innervation are supplied through the apical foramen, a narrow opening at the apical end of each dental root. This anatomical configuration often makes relatively minor inflammation result in severe pain and tissue necrosis, by strictly limiting tissue swelling leading to an increased internal pressure. If tissue vitality is preserved then damaged tissue can be recovered by the regenerative abilities of DP. DP regenerates by recruiting the same cell types involved in homeostasis, and no new cell type that is unique to injury has been observed (1-3). Although DP tissue repair can be promoted by various dental treatments such as pulp capping, few interventions harness its extensive regenerative properties. Currently, the composition of DP at the cellular and molecular levels, as well as the mechanisms by which DP resident cells maintain homeostasis, are not well understood.

Several studies have reported interactions among cells in DP, with special attention given to mesenchymal stem cells (MSCs) and their niches (4,5). It is now accepted that DP possesses two main MSCs niches, perivascular and neural, which participate in tissue development and regeneration (6). Within these niches, mechanisms of self-regulation have been discovered. However, it is not clear how other resident and transient cells interact with these niches or participate in the regulation of overall tissue homeostasis.

Single-cell transcriptomics allows the quantification of gene expression in individual cells in a tissue sample. Researchers have used this technology to generate transcriptomes of developing teeth in mice (7,8). Recent studies of human DP have identified cells involved in tissue development (2), delineated major cellular components of adult DP (9), and described enriched cell types under carious lesion conditions (10).

Here, we sought to characterize DP tissue at the cellular and molecular levels by integrating the data from all publicly available healthy human DP single-cell transcriptomics datasets. As a reference, we also integrated data from five different human tissues in healthy adults. We hypothesized that the gene expression patterns in DP cells would be distinct from those in the corresponding cell types in reference tissues. We further reasoned that, if such unique phenotypes were observed, they would help shed light on the biological processes responsible for DP tissue homeostasis. Such insight may further inform the development of cellular and molecular-based treatment strategies in dentistry.

## Methods

### Dataset selection

FASTQ files of single-cell RNA sequencing (scRNA-seq) data were retrieved from published studies (2,9-13). The data were selected based on healthy human tissue. All selected datasets were generated by the Chromium single-cell gene expression platform from 10X Genomics 5’ V1 and 3’ (V 2 or V3) to avoid other confounding factors and technical biases. In total, we included nineteen datasets from six different tissues; nine DP (GSE185222, GSE161267, and GSE146123), two bone marrow (BM) (PRJEB37166), two adipose (ADP) (GSE155960), two lung (PRJEB52292), two skin (PRJNA754272) and two peripheral blood mononuclear cells (PBMC). The two PBMC data were acquired from publicly available datasets from 10X Genomics (11) (**Supplementary Table 2**).

### Data pre-processing and quality control

Raw FASTQ files of each dataset were processed with 10x Genomics Cell Ranger version 7.0.0 and mapped against the human reference genome GRCh38 version 2020-A (GENCODE v32/Ensembl 98). Count matrices were then processed with the Seurat package (version 4.1.1) (14) in R (version 4.2.0). Quality control was performed by excluding the cells expressing mitochondrial genes higher than 10% or those expressing lower than 200 genes or higher than 4,000 genes. Cells that expressed more than 3,500 genes were filtered out for the integration of nine dental pulp datasets. Genes that were detected in fewer than three cells were discarded. Data were normalized by the log normalization method and scaled using the NormalizeData and ScaleData functions in Seurat (14) to yield a total count per cell of 10,000.

### Data integration and visualization

Data was integrated by the Seurat integration pipeline (15) using 2,000 high variable gene features across all datasets. The top thirty components from principal component analysis (PCA) were used to calculate a Uniform Manifold Approximation and Projection (UMAP) dimensionality reduction, after which clusters were identified using Louvain algorithm (16) based on a shared nearest neighbor (SNN) in the FindCluster function. Each cluster was manually annotated using cell type markers from the literature and curated markers from CellTypist (17).

### Quantifying transcriptomic expression of clusters and pseudo-bulk data

Pseudo-bulk PCA was performed using average gene expression across all cells within a cluster. The union of the top 500 genes having the highest absolute values for PC1 and PC2 were retrieved. The average expressions of these genes were transformed into Z-scores and subsequently used to construct a heatmap comparing gene expression profiles in DP with those of the other five tissues. The PCA plots and heatmaps were constructed using ggplot2 (Wickham, 2016) and ComplexHeatmap R packages (Gu et al., 2016)

### Differential gene expression and gene ontology (GO) analysis

Fibroblast clusters were subset from the integrated dataset of the six tissues. Differential gene expression (DE) analysis of fibroblasts in DP and in other tissues was performed by the non-parametric Wilcoxon rank sum test using the FindMarkers function in the Seurat pipeline with gene expression levels before integration. The DE results were visualized by a Volcano plot using the scmisc R package (version 0.8.0). GO enrichment analysis for Biological Processes ontology was performed with the list of DE genes with FDR<0.05, separating up-regulated and down-regulated genes. This analysis was done using the goana function implemented in the limma package (18) and wrapped for single cell analysis in the scmisc package.

### Cell-Cell interaction analysis

Interactions between cells were predicted based on the expression levels of ligands and their known interacting partners by CellPhoneDB (19). A heatmap showing potential interactions between cell types was constructed using the default visualizing function of CellPhoneDB. Dot plots showing the interacting pairs and the levels ligand-receptors expression were performed using the scmisc package. Significant means of ligand-receptor interactions from fibroblast clusters were summed to obtain a total interaction score between fibroblast and other cell types. The top 25 interactions were selected to create an adjacency matrix using the igraph R package (20) which was subsequently used to plot the chord diagrams with the chordDiagram (21) package in R.

## Results

### DP resident fibroblasts exhibit unique gene expression profiles

The cell composition of and transcriptomic signatures within DP tissues were investigated by integrating nine DP single-cell RNA sequencing datasets with ten datasets from five reference tissues: PBMC, BM, ADP, LUNG and SKIN. We selected these tissues because they contain representatives of the major cell types populating the DP (6,22). After applying all the QC filters, this resulted in a dataset with 109,554 cells, and 33,401 genes, including 48,659 cells from DP and 60,895 from other tissues (**Fig 1A**). Clustering identified nineteen distinct cell clusters which were further annotated by cell type (**Fig 1B)** (**Supplementary Fig 1A**) using established markers from the literature along with CellTypist encyclopedia (23) (**Supplementary Fig 1B**). Overall, fibroblasts which were identified by the expression of markers COL1A1, COL1A2, COL3A1, DCN, and CXCL14 (**Fig 1C**) were the most abundant cell type. We found a very small cluster of MKI67+ proliferating fibroblast (pro-fibroblast) (Cluster18) that was present only in skin tissue (**Fig 1D**). Schwann cells (Schw), which consisted of myelinated Schw (MBP+) and non-myelinated Schw (GFRA3+) were identified by the expression of SOX10 (24-26) (**Supplementary Fig 2**). DP was enriched in both fibroblasts and Schw clusters (Fig 1D). In addition, a small cluster of odontoblasts were identified by the expression of DMP1 (**Fig 1B**) (**Supplementary Fig 3**). As expected, odontoblasts were observed only in DP tissues. Other cells identified included immune cells (cluster 8, 13, 15, 1, and 14), endothelial cells (clusters 6 and 12), epithelial cells and keratinocytes (cluster 16). Finally, two clusters (5 and 10) with markers compatible with perivascular cells (CSPG4) (27), including vascular smooth muscle cells (ACTA2, TAGLN, and TPM2) (12,28), and mesenchymal stem cells (ACTA2, FRZB, NOTCH3, and MYH11) (9,27) were annotated as MSC-like cells (**Supplementary Fig 4**).

**Figure 1.**
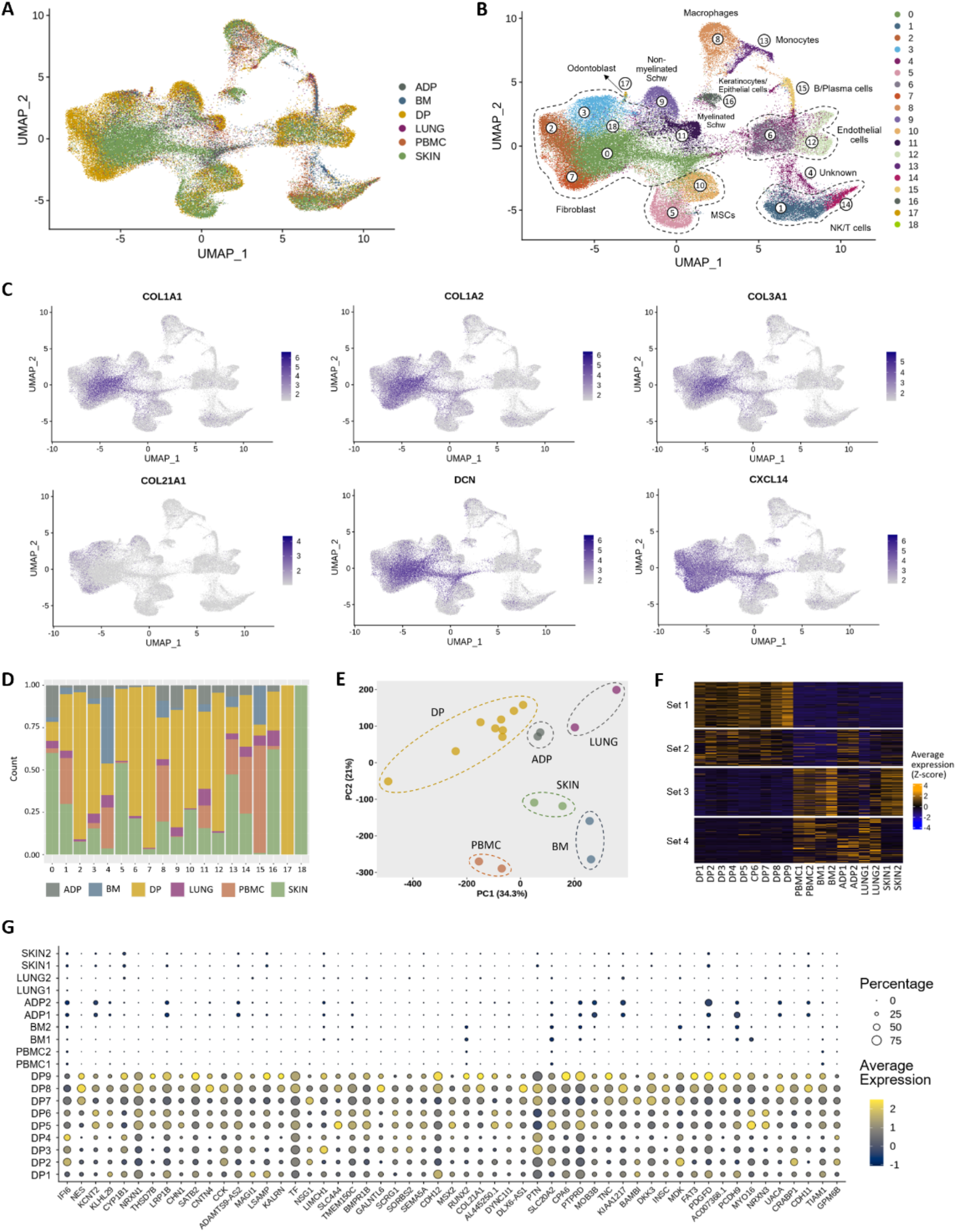
scRNA-seq analysis of human dental pulp and reference tissues from healthy adults. (**A**) UMAP visualization of all integrated tissues, cell clusters are colored by tissue of origin. (**B**) UMAP visualization of unbiased cluster classification identifying 19 cell clusters with cell type annotation. (**C**) Feature plots of fibroblast markers identifying the cell population within clusters 0, 2, 3, 7 and 18. (**D**) Bar plot of the fraction of cells that compose each cluster shown in **C**, colored by tissue of origin. (**E**) Principal component analysis plot of fibroblasts, performed using average gene expression for each sample. (**F**) Heatmap of the top 500 differentially expressed genes that contributed to PC1 and PC2 in **E**, set 1 shows upregulated genes in DP fibroblasts. (**G**) Dot plot of curated genes of interest retrieved from heatmap set 1, the full list of genes is provided in **Supplementary Table 1**.

To identify tissue-specific features, we performed PCA for each cell cluster in **Fig 1B** using average expression profiles (pseudo-bulk) for each sample. The corresponding plots showing the pseudo-bulk sample distribution in the first 2 principal components (PCs) of all clusters is shown in **Supplementary Fig 5**. The PCA plots indicated that DP fibroblasts (Cluster 0, 2, 3, and 7) and Schwann cells (Cluster 9 and 11) were well separated from those of other reference tissues, suggesting the existence of DT specific markers driving this separation (**Supplementary Fig 5)**.

In order to understand the gene expression signatures unique to DP fibroblasts, we performed PCA on pseudo-bulk signatures using cells in the fibroblast clusters (0, 2, 3 and 7). For this analysis, proliferating fibroblasts (cluster 18) were not included, as they were found only in skin. The first two components (PC1 and PC2) showed separation between fibroblasts from DP compared to fibroblasts from other tissues (**Fig1E**). Then, the top 500 genes contributing to PC1 and PC2 were obtained and used to construct a heatmap (**Fig1F**). The heatmap highlighted a group of genes (set 1) as being upregulated in DP fibroblasts while downregulated in fibroblasts of other reference tissues. We further curated genes from the heatmap set 1 to construct a dot plot with genes of interest showing the highest average expression in DP (**Fig 1G**) (a full list of genes retrieved from heatmap set 1 is provided in Supplementary Table 1). Taken together, our analyses suggest that DP fibroblasts have a unique gene expression profile when compared to the fibroblasts of the five reference tissues.

### Heparin-binding growth factors pleiotrophin (PTN) and midkine (MDK) are highly expressed in DP resident fibroblasts

To gain insight into DP fibroblast specific genes at single-cell resolution, we next compared the differential gene expression between DP fibroblasts and those from reference tissues. The volcano plot in **Fig 2A** highlights genes with large fold changes that are also statistically significant. The eight most elevated genes in DP fibroblasts were PTN, TF, PTPRD, NRXN1, CDH12, CCK, CPA6, and MDK (**Fig 2A**), consistent with the genes retrieved from PCA pseudo-bulk analysis shown in **Fig 1G**. We next performed gene set enrichment analysis to investigate which gene ontology (GO) terms were under- or over-represented in these genes. The top GO terms in the Biological Process ontology were: “nervous system development”, “cell morphogenesis”, “neurogenesis”, “system development” and “cell differentiation” (**Fig 2B**).

**Figure 2.**
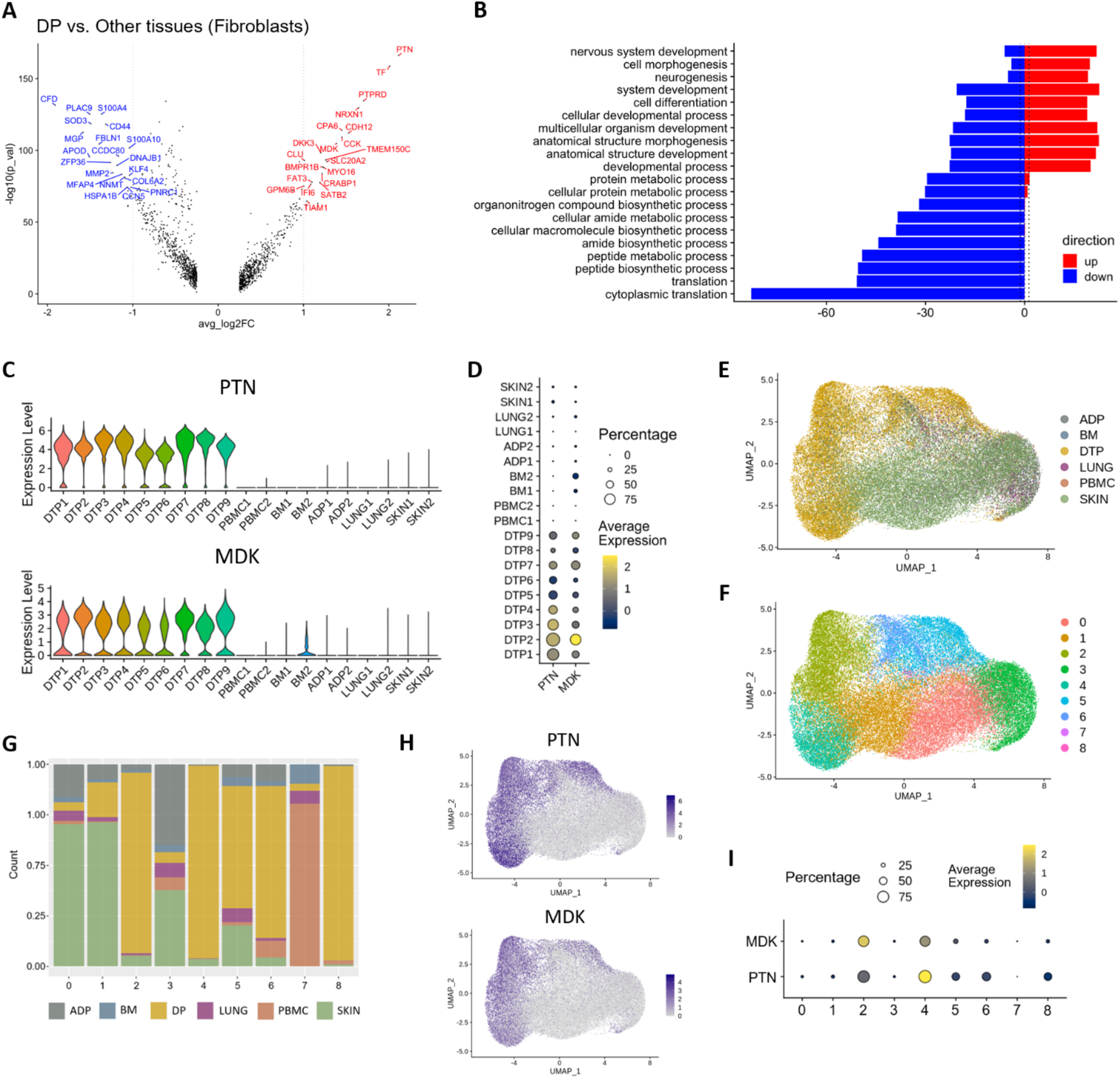
Dental pulp fibroblasts versus those from reference tissues. (**A**) PTN and MDK are differentially expressed genes in DP fibroblast versus reference tissue fibroblasts. (**B**) Gene ontology enrichment analysis using the DE genes from **A** identified significantly upregulated Biological Processes in DP fibroblasts. (**C**) Violin plots of PTN (upper) and MDK (lower) expression levels in fibroblasts of all tissues. (**D**) Dot plot showing percentage and average expression of PTN and MDK of fibroblasts of all tissues. (**E**) UMAP visualization of fibroblasts from all tissues integrated and colored by tissue of origin. (**F**) Unbiased classification identifying 9 clusters of fibroblast sup-populations. (**G**) Bar plot of the fraction of cells that compose each cluster shown in **F**, colored by tissue of origin. (**H**) Expression of PTN (upper) and MDK (lower) projected onto the UMAP plot of integrated fibroblasts. (**I**) Dot plot showing percentage and average expression of PTN and MDK by clusters of fibroblasts in **F**.

Within the DE genes in **Fig 2A**, PTN–the most upregulated gene in DP fibroblasts–and MDK, are paralogs coding for Pleiotrophin and Midkine, respectively. PTN and MDK are members of a distinct family of secreted heparin-binding growth factors that have been implicated in various biological processes, from development to inflammation to tumorigenesis (29). We found that PTN and MDK were highly expressed in all nine DP datasets, while their levels were extremely low in all ten datasets from the other five reference tissues (**Fig 2C-D**).

To further investigate the unique contribution of PTN and MDK to DP fibroblasts, fibroblast clusters from all six tissues were independently integrated, and UMAP coordinates and clusters were recalculated, resulting in nine subpopulations (**Fig 2E-F**). Fibroblast clusters 2, 4, 5, 6 and 8 were enriched significantly in DP tissues (**Fig 2G**). As expected, PTN and MDK were expressed only in these DP-resident fibroblast clusters (**Fig 2H-I**), confirming that PTN and MDK are the key genes distinguishing the DP fibroblasts from the reference tissue fibroblasts.

### Analysis of DP cell populations and PTN- and MDK-mediated intercellular communication networks

To understand the biological relevance of PTN and MDK in the DP environment, we next ignored the reference tissues and turned our attention exclusively to the DP datasets. Data integration of all nine healthy DP datasets retrieved from three different studies (Supplementary Table 2) was performed. Unbiased clustering predicted nineteen different clusters, which were further grouped into eleven cell types (**Fig 3A**) using known markers indicated in **Fig 3B**. Our results revealed that fibroblasts were the largest cell population in the dental pulp (43.92%), followed by endothelial cells (18.05%), MSCs (13.79%) and Schwann cells (12.39%) when combining non-myelinated and myelinated cells. While B cells and erythrocytes were the smallest cell cluster (0.4%) (**Fig 3C**). As expected, elevation of PTN and MDK expression was found in clusters annotated as fibroblasts (**Fig 3D**).

**Figure 3.**
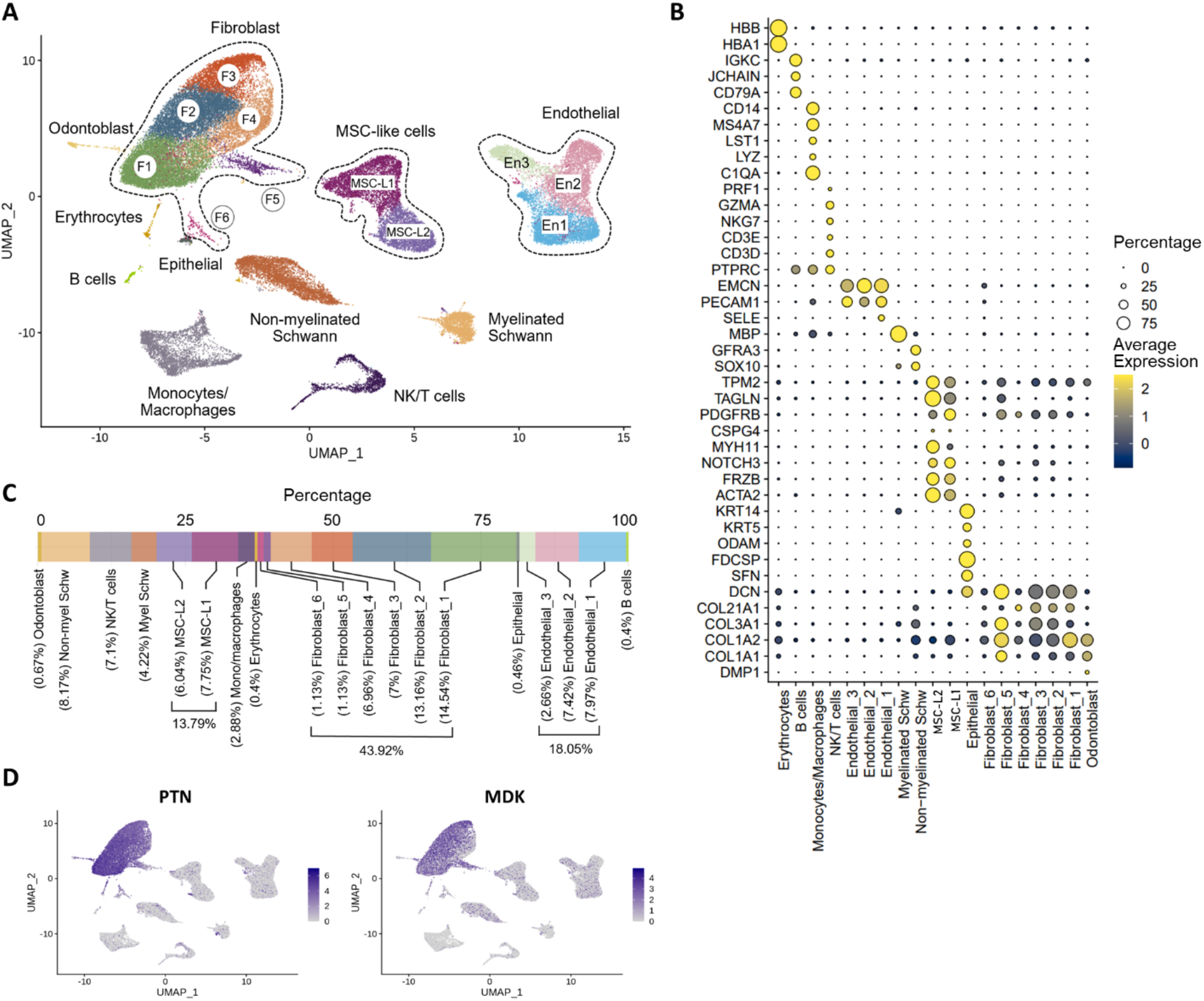
ScRNA-seq analysis of integrated human dental pulp tissues. (**A**) UMAP visualization of integrated DP datasets. (**B**) Expression levels of known cell type markers used for the cluster annotation in **A**. (**C**) Percentage of cell types that compose the healthy adult human DP. (**D**) Expression of PTN (left) and MDK (right) projected onto the UMAP plot of integrated dental pulp tissues, fibroblasts highly express PTN and MDK.

Since PTN and MDK are secreted heparin-binding molecules, we sought to investigate potential target cells based on the mRNA expression of their known receptors using CellPhoneDB (19). Our analysis revealed a high number of interactions between fibroblasts and non-myelinated Schwann cells, MSCs, endothelial cells and odontoblasts (**Fig 4A**). To illustrate specific ligand-receptor pairs contributing to these interactions between fibroblasts and other cell types, chord diagrams were constructed using the top twenty-five interacting pairs (**Fig 4B**). It is shown that PTN and MDK were the top molecules mediating major communication between fibroblasts and non-myelinated Schwann cells, MSCs and odontoblasts, but not endothelial cells (**Fig 4B, Supplementary Fig 6**). In addition, even though fibroblast interacted with several DP cells via PTN and MDK, the pattern of receptor usage varied depending on the cell type (**Fig 4B, Supplementary Fig 6**).

**Figure 4.**
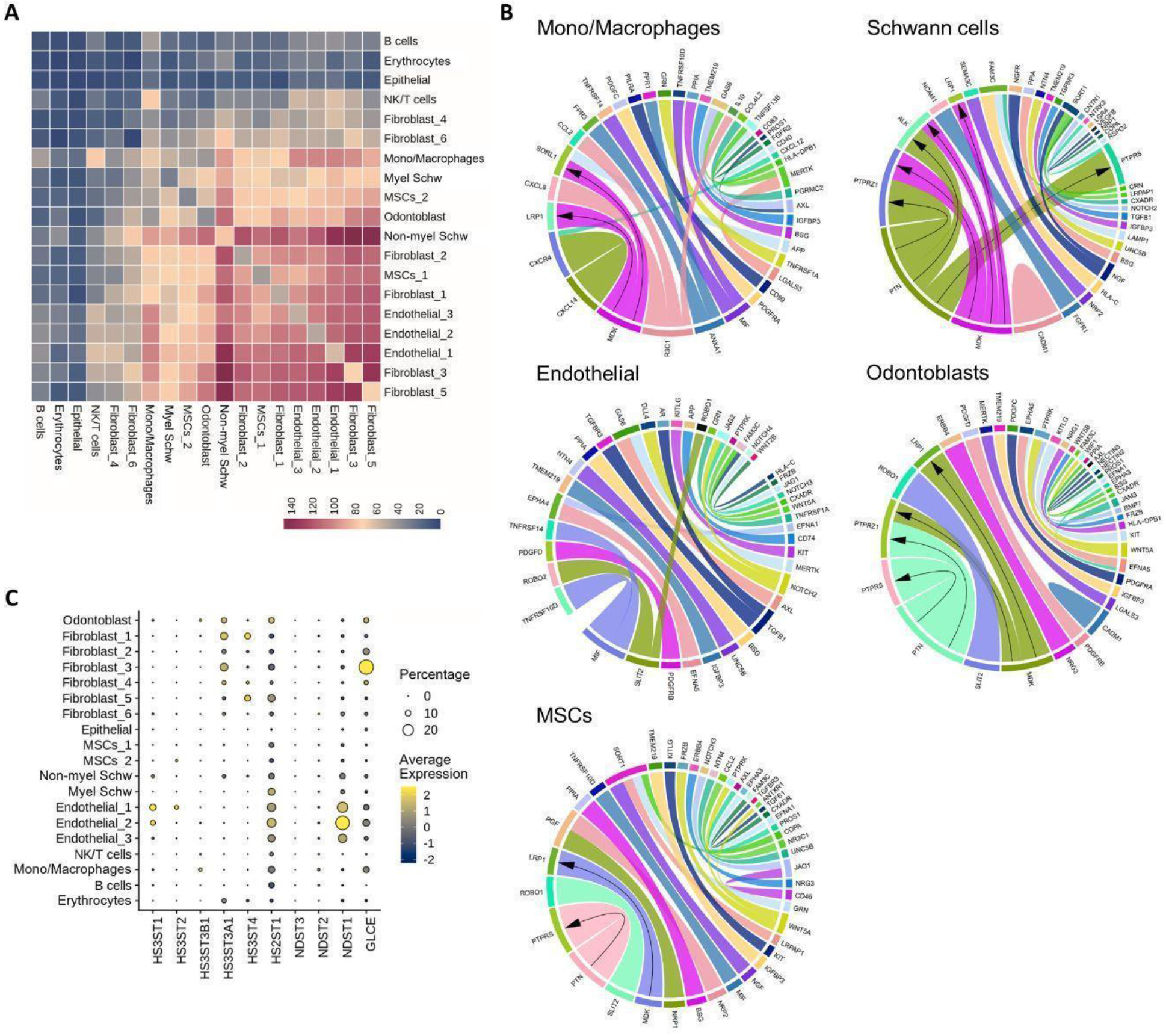
CellPhoneDB analysis predicted PTN and MDK interactions in the dental pulp. (**A**) Heatmap showing the total number of statistically significant predictions of cell-cell interactions. (**B**) Circus plot/chord plot of the top 25 ligand-receptor pairs between fibroblasts and their corresponding cell types. Line thickness represents an interaction score obtained from fibroblast clusters involved in the interaction. Arrows indicate PTN and MDK on fibroblasts and their potentially interacting partners on particular cell types. (**C**) Dot plot showing the percentage and expression levels of genes involved in the biosynthesis of heparan sulfate molecules.

PTN and MDK bind both heparin and heparan sulfate (HS). These are linear polysaccharides attached to cell surfaces by one or more anchoring proteins. In order to further support the role of PTN- and MDK-mediated communication between DP fibroblasts and various other DP cells, we next investigated the expression levels of genes encoding major HS anchor proteins. We found BGN, GPC3, SDC2 were expressed on cells that also expressed high levels of receptors for PTN and MDK (**Supplementary Fig 7**). In addition, we determined the expression of genes involved in the biosynthesis of HS itself (**Fig 4C**). Taken together, these results support a model where PTN and MDK are produced by DP resident fibroblasts and exert a paracrine effect on several types of DP cells, including Schwann cells, endothelial cells, and odontoblasts.

## Discussion

In order to characterize the cellular and molecular machinery of DP tissue, we integrated all publicly available single-cell gene expression data from healthy human DP and compared the expression profiles with several well-studied reference tissues. We hypothesized that, due to the unique functional constraints on DP cells, their gene expression patterns would be distinct from those of other tissues. Our analysis revealed that DP tissue indeed exhibited a unique gene expression profile in comparison to reference tissues and that DP resident fibroblasts are the source of these differences. Other cellular components of the tissues studied here, such as endothelial cells and immune cells, did not show significant differences, suggesting a common behavior across those tissues. While fibroblasts are found in all tissues, their function is dependent on local environment and context to support other resident cells and engage in tissue remodeling when necessary (30,31). In our study, we evaluated data obtained from healthy adult DP tissues, and assumed a context of steady state and homeostasis. A total of six subpopulations of fibroblasts were identified, revealing a previously unappreciated diversity, suggesting that each distinct fibroblast subtype within DP may be involved in distinct functional roles. Characterization of these subpopulations is now necessary to determine these functional roles, and particular attention should be placed on the identification of non-overlapping functions, as these may be used to develop new therapeutics.

Studies on single-cell analysis of human DP are few, and their methodologies differ both in sequencing platform and chemistry. To minimize batch effects in our analysis we selected only studies that used equivalent and most current methodology, ignoring some recent studies that used different platforms. However, our results largely agree with previously published data, regardless of their methodology. Yin et al identified similar cell types as were observed in our analysis; a cluster labeled “pulp cells”–which can be regarded as fibroblasts given their expressed markers–showed an active communication network with other cells in the DP (32). MSCs were found to be the main cluster interacting with fibroblasts; however, we show here that, in addition to MSCs, fibroblasts have significant interactions with Schwann cells, endothelial cells and odontoblasts. The integrative approach used here, that included multiple datasets from the literature, yielded a greater number of cells than any individual study; our integrated dataset could capture additional significant interactions in the communication networks within cell clusters.

Our analysis identified several functional genes contributing to the unique expression profile of DP-resident fibroblasts, and involved in tissue growth: Transferrin (TF), reported to support tooth morphogenesis and dental cell differentiation (33); Cadherin 12 (CDH12), which plays an important role in calcium-dependent cell adhesion and has been implicated in oral tumors (34); Cholecystokinin (CCK), which regulates calcium transport in teeth (35). Notably, genes related to nervous system development and neurogenesis were differentially expressed, a finding reiterated by the GO analysis. Receptor-type tyrosine-protein phosphatase delta (PTPRD), identified as a critical growth suppressor in the central nervous system (CNS) (36); Neurexin 1 (NRXN1), a receptor found in CNS synapses that plays an important role in neural development (37); Carboxypeptidase A6 (CPA6), involved in the processing of neuropeptides and reported to be upregulated in DP (38). Taken together, these findings are consistent with the ectomesenchymal origin of DP. The embryological development of craniofacial connective tissues involves neural crest-derived cells that acquired mesenchyme features, and thus are termed ectomesenchymal cells (39). This neural component of DP tissues remains relevant after development. In adult tissue, peripheral axons comprise 40% of DP volume (40), are the basis for pain processing (41), and participate in immune response (42,43). Although neurons cannot be isolated by the tissue digestion techniques applied in the present work, we were able to show that glial cells alone, myelinated and non-myelinated Schwann cells, comprise 12.39% of the DP cell population. It is possible that the mechanisms for suppression of cell proliferation *in vivo*, and for the overall regulation of cellular turnover, emerge from DP-resident fibroblasts and glial cells through paracrine regulation.

Among the differentially expressed genes in DP-resident fibroblasts PTN appear as the top significant and MDK is its highly conserved paralogue. Both PTN and MDK code for biochemically similar proteins that share 45% identity in amino acid sequence and exhibit strong binding to sulfated glycosaminoglycans, including heparin, heparan sulfate and chondroitin sulfate (29,30,44). These heparin-binding growth factors are involved in several important molecular processes, among which neural development and tumorigenesis have been well studied (29). Recently, Zhang and co-workers examined the role of PTN in DP stem cell maintenance, and, using RNA interference in cultured DP, showed that PTN protects DP stem cells from senescence (45). MDK is essential in tooth development, participating in epithelial-mesenchymal interactions, where it is secreted by cells of the dental papilla mesenchyme and captured by epithelial cells (46). This mechanism was further elucidated in a later report, using dental papilla cells *in vitro*, where MDK could promote cell differentiation into odontoblast-like cells via the modulation of *dspp* (47). The role of MDK in the adult healthy DP tissue homeostasis is largely unknown, high expression of MDK has been previously identified by microarray screening, but expression levels dropped dramatically when compared to caries conditions (48). Together, these studies indicate that MDK is involved in cell proliferation and differentiation but the circumstances leading to the presence or absence of MDK are unclear.

Cell-cell communication predictions indicated that among the top 25 significant pairs, PTN/PTPRZ1 may be especially relevant to understanding DP homeostasis. Upon receptor binding, PTN inactivates the intrinsic tyrosine phosphatase activity of PTPRZ1 (49,50), affecting several downstream targets that are highly regulated by phosphorylation. These include β-catenin (50) and Fyn kinase (51), which are essential to cell proliferation and differentiation, to cell adhesion and cell motility, and regulation of apoptosis. Similarly, MDK binding to PTPRZ1 is predicted to have analogous effects given their homology (29,52). It is known that primary cultures of DP stem cells from explanted adult DP shows a high proliferation rate *in vitro* (53), however, this is not observed in vivo where cell mitosis is at times undetectable (54-58). Our analysis revealed for the first time that the mechanisms for tissue homeostasis, through which DP controls its extraordinary proliferative and differentiation abilities, may involve PTN/PTPRZ1 binding or their homologues.

Growth factors extracted from human platelets are currently used for promoting dental pulp regeneration (59). As life expectancy has increased, thanks to the advancement of science and technology, the maintenance, repair, and regeneration of dental tissue has become ever more important. However, there has been little research into the biochemical pathways specific to DP that regulate tissue growth or repair. The identification of PTN and MDK are a first step in this direction. The extensive receptors for these proteins on Schwann cells, along with the other factors involved in CNS development or regulation, imply extensive crosstalk between fibroblasts and nerve cells in the regulation of calcium and other essential components of teeth. Synthetic molecules that are able to target DP accurately and specifically are currently lacking. This study serves as a stepping stone for tailored molecular dental treatments. Design and synthesis of DP-targeting molecules will be an important next step. Systematic discovery and straightforward delivery of such molecules would provide effective materials for durable dental tissues and greatly expand the arsenal of potential dental treatments.

## Acknowledgements

The authors thank Jantarika Kumar Arora for single-cell analysis consultancy. We would like to acknowledge the support from the FrontierLab exchange program from Osaka University in enabling this study.

## Authors contribution

Conceptualization and study design: N.J., G.L.A., D.M.S., Data analysis: N.J., M.J.L.L., D.D., Preparation of the first draft manuscript N.J., G.L.A., D.M.S., Review and editing: S.P., P.M., J.S., S.I.

All authors have reviewed and approved the submitted manuscript

## Code availability

Code to reproduce the analyses described in this manuscript can be accessed via: https://github.com/NatnichaJira/DentalPulp

## SUPPLEMENTARY INFORMATION

**Supplementary figure 1.**
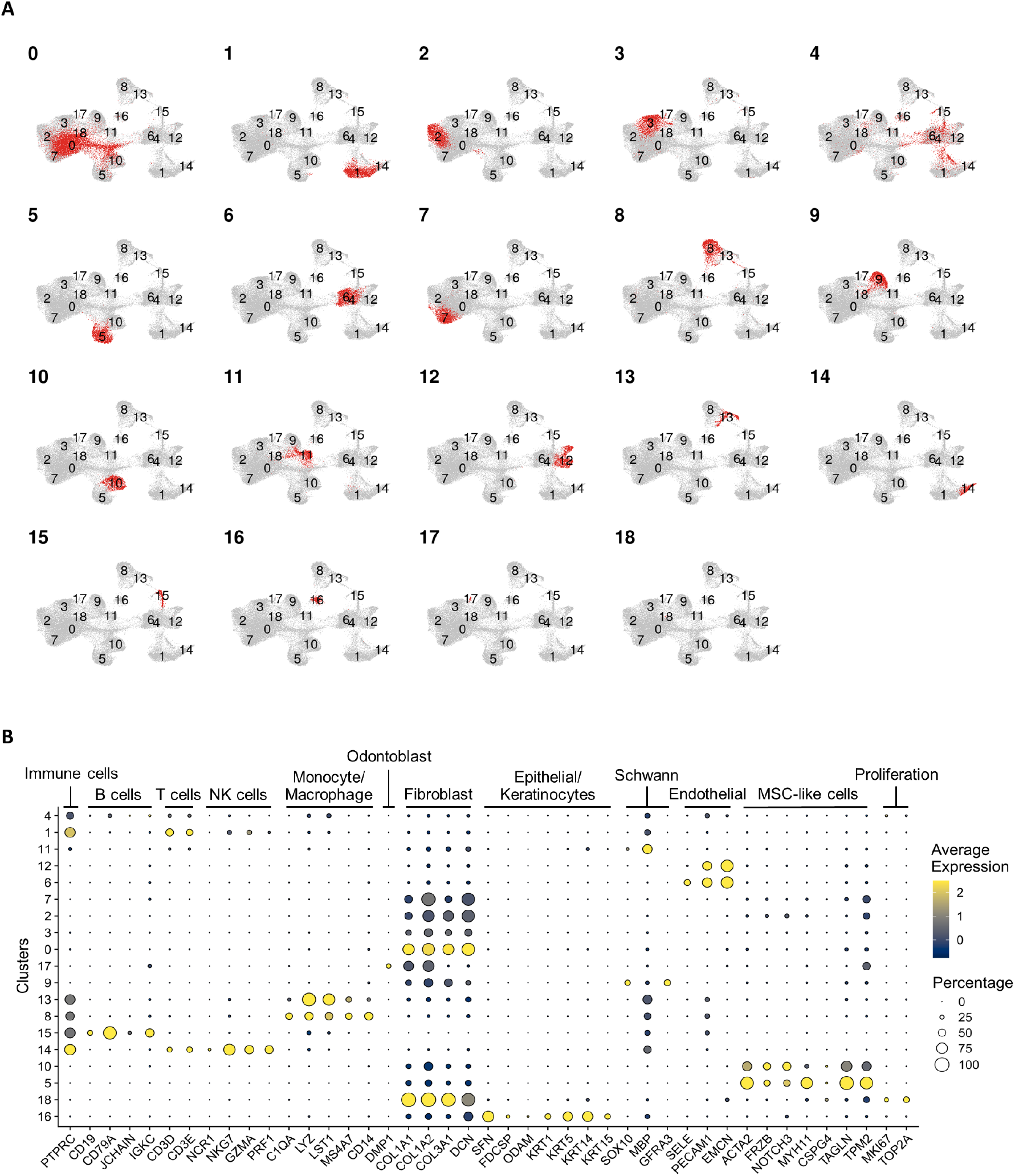
Cell type annotation of an integrated data of 6 different tissues. (**A**) UMAP plots showing the location of each cell types. (**B**) Gene markers used to annotate cell type clusters.

**Supplementary figure 2.**
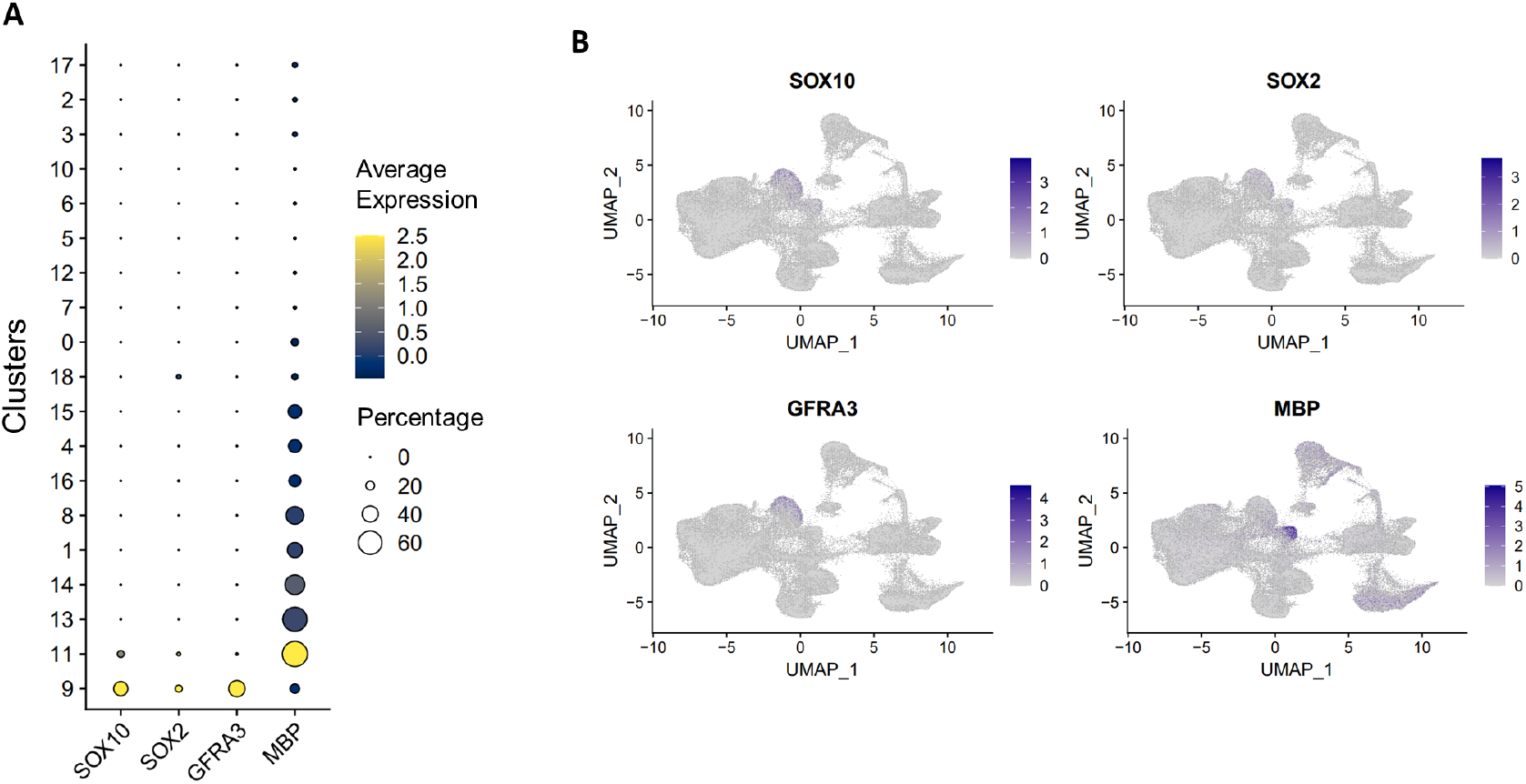
Annotation of myelinated and non-myelinated Schwann cells. (**A**) Dot plots showing the average and percentage of marker gene expressions. (**B**) Expression level of marker genes on the UMAP plot in Figure 1A showing that cluster 9 was non-myelinated Swann cells while cluster 11 was myelinated Schwann cells.

**Supplementary figure 3.**
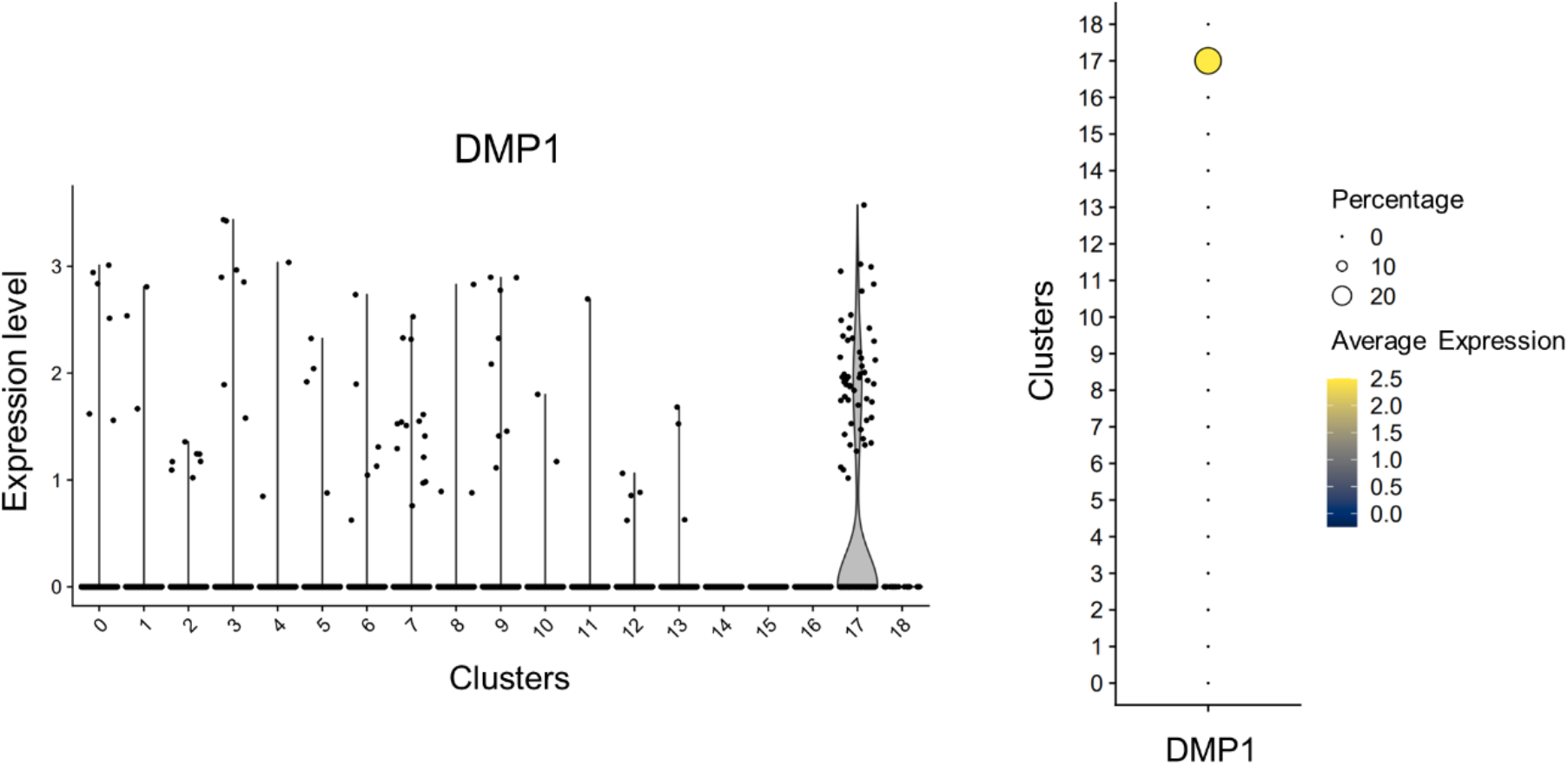
DMP1 expression in cluster 17 (Odontoblast).

**Supplementary figure 4.**
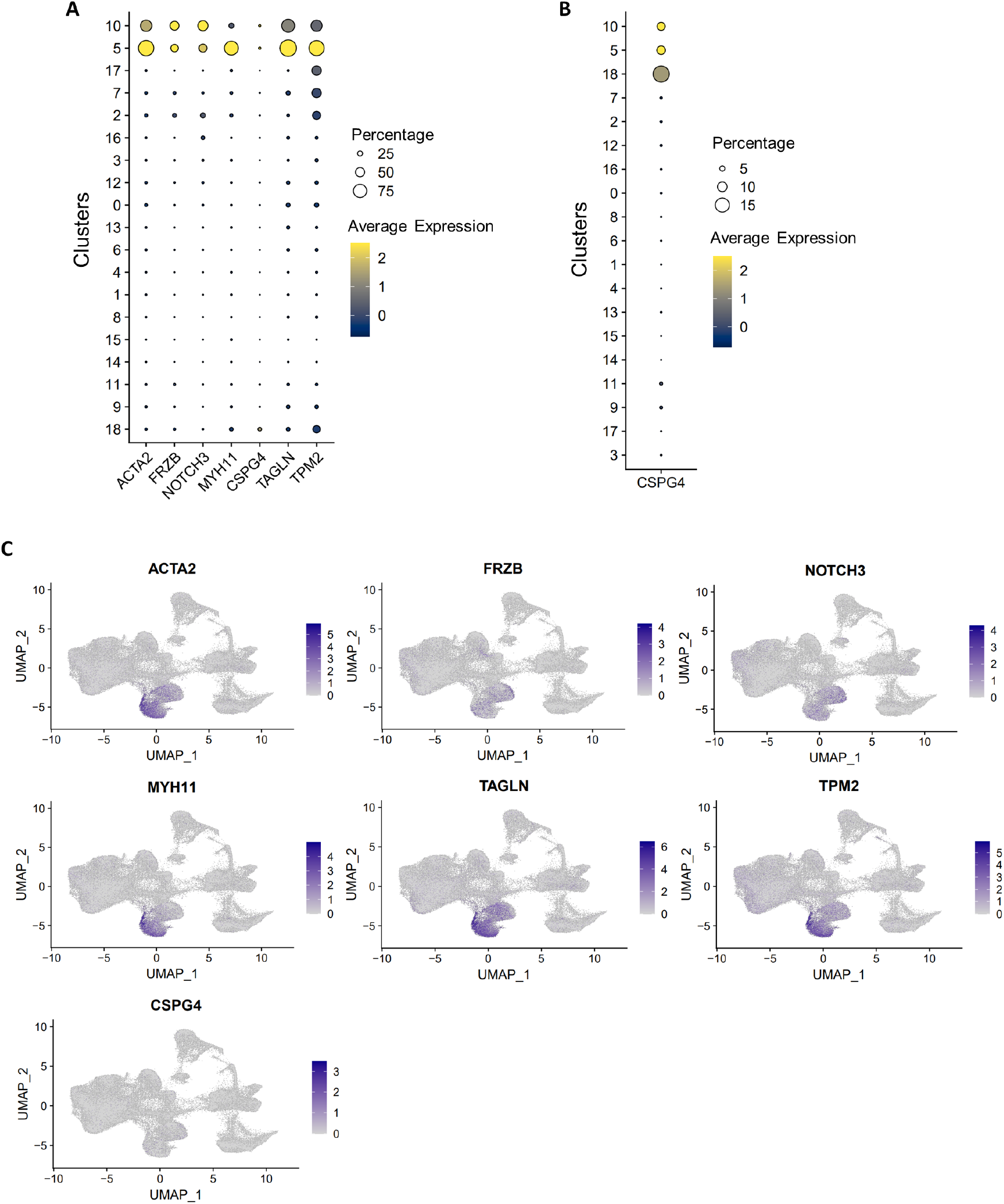
Annotation of MSC-like cells. (**A-B**) Dot plots showing the average and percentage of marker gene expressions. (**C**) Expression level of marker genes on the UMAP plot in Figure showing that cluster 5 and 10 were MSC-like cells.

**Supplementary figure 5.**
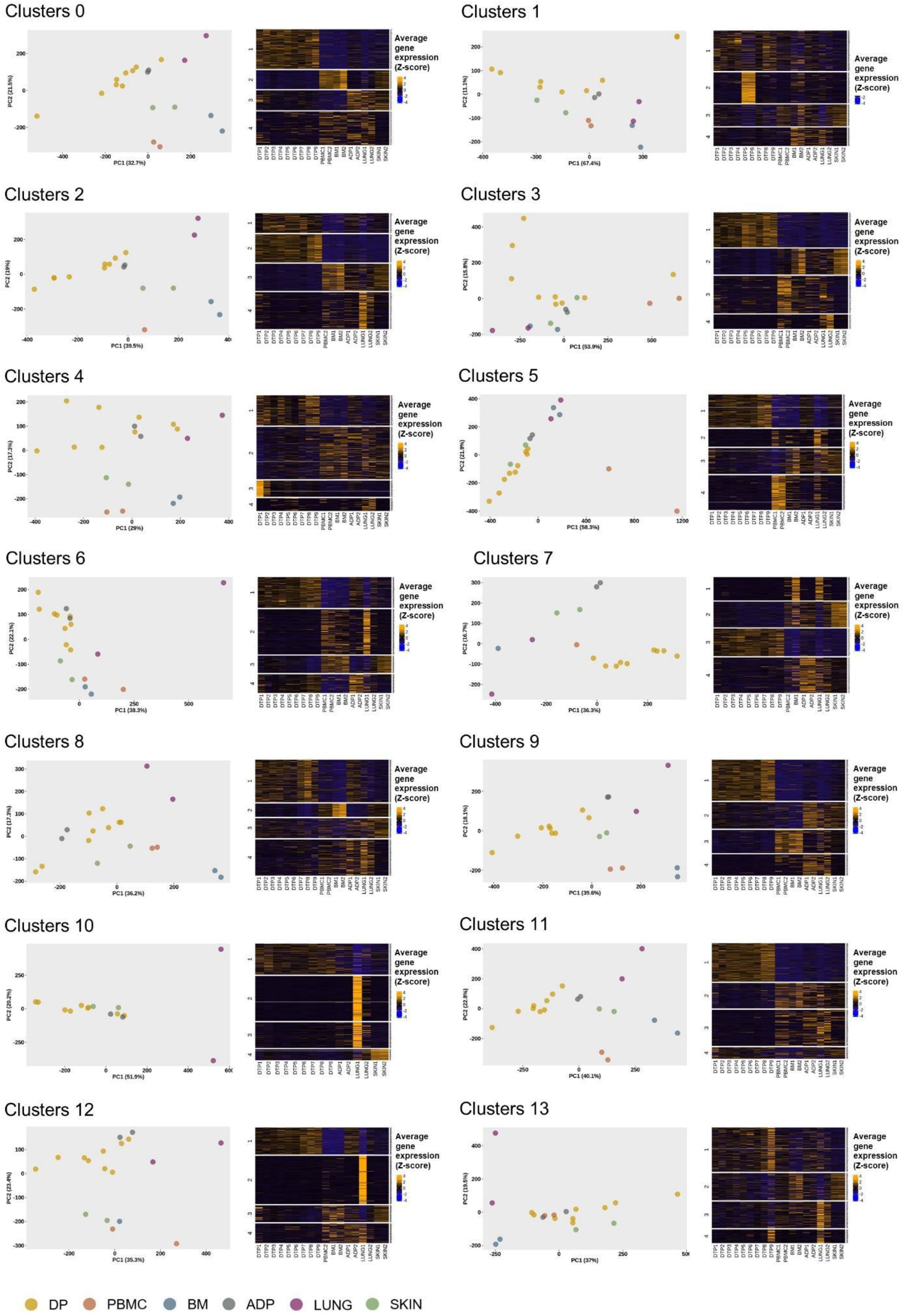
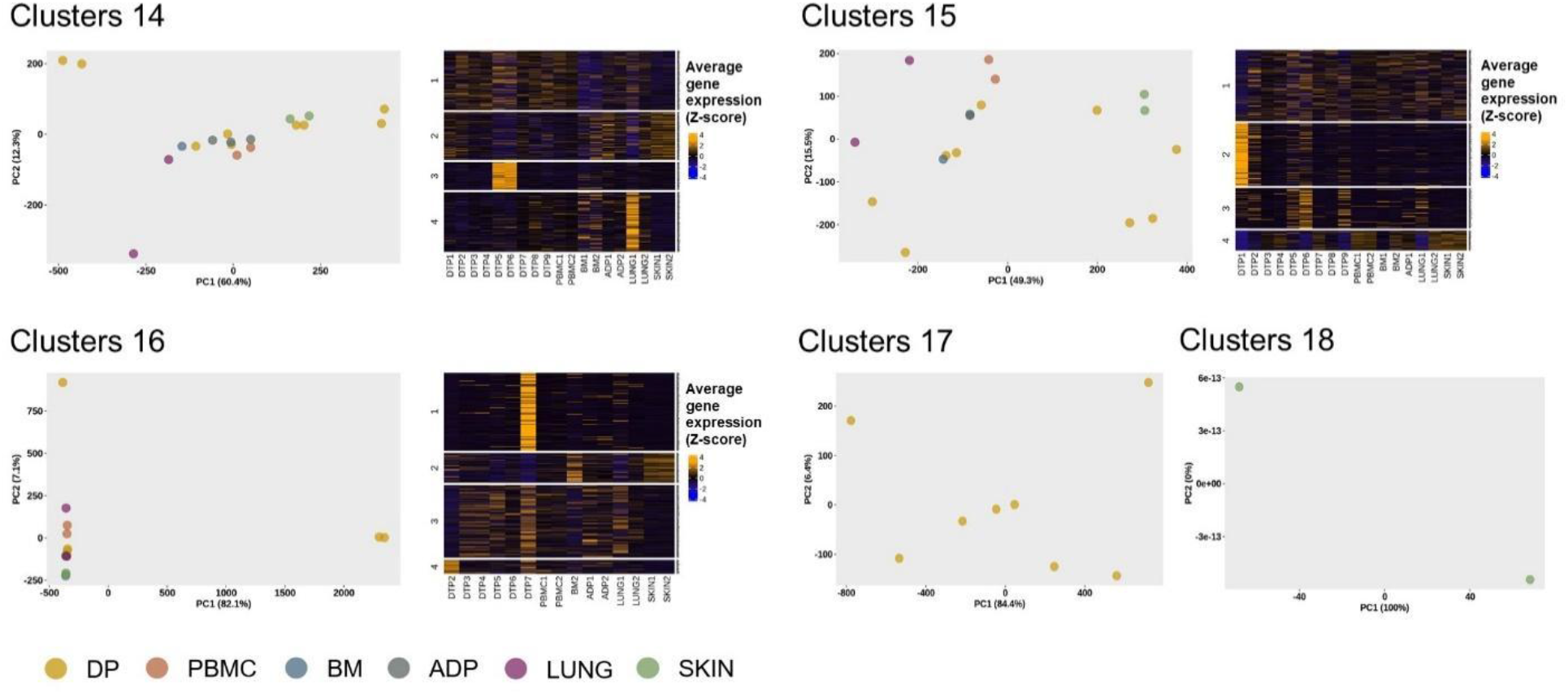
Pseudo-bulk PCA and heatmap of all clusters showed in Figure 1B. PCA were performed using average gene expression profile for each samples. The top 500 genes contributing to PC1 and PC2 were retrieved and used to construct a heatmap.

**Supplementary figure 6.**
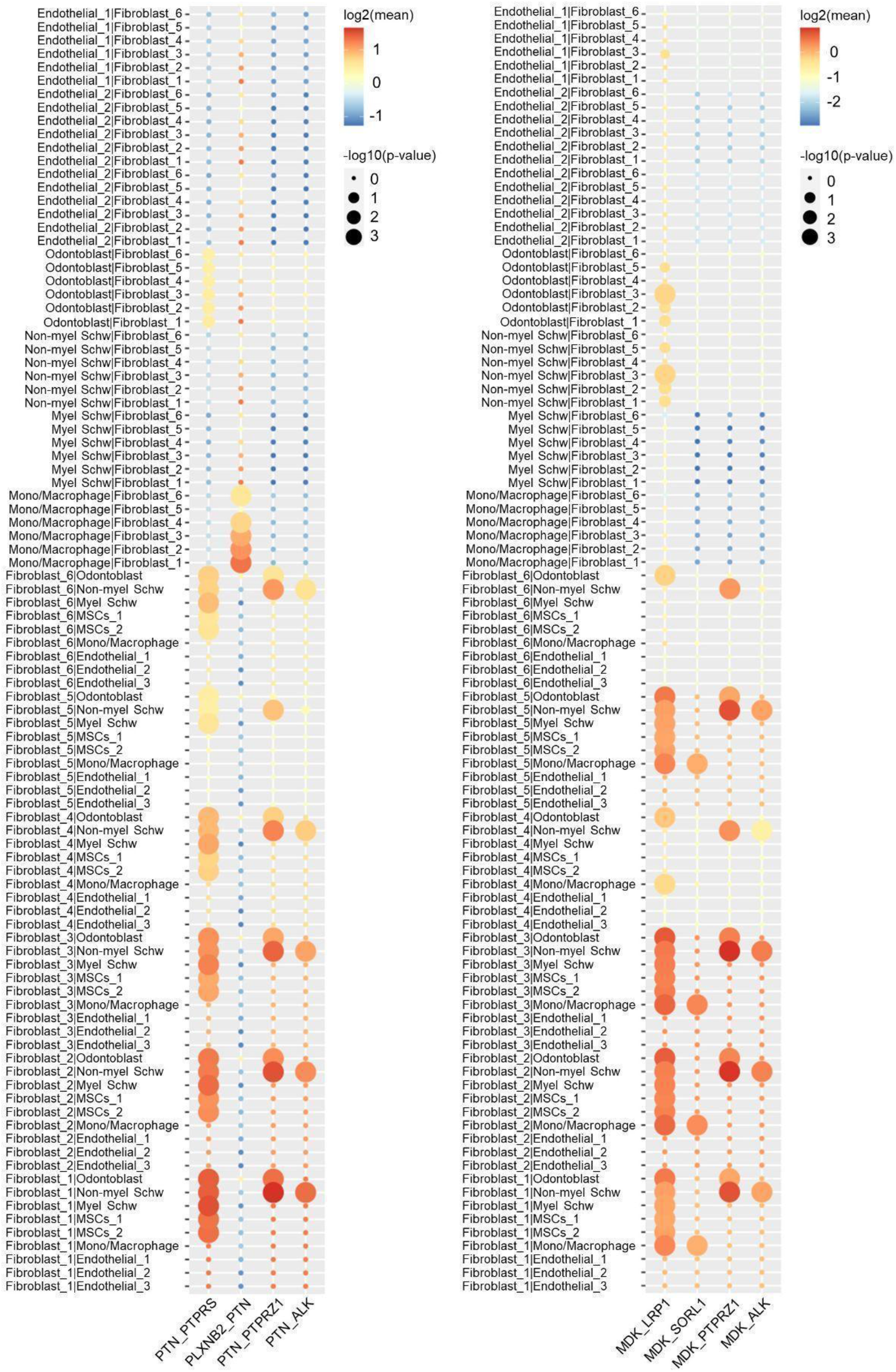
Mean and p-value of average expression levels of PTN and MDK and their interacting partners between fibroblasts and other cell types

**Supplementary figure 7.**
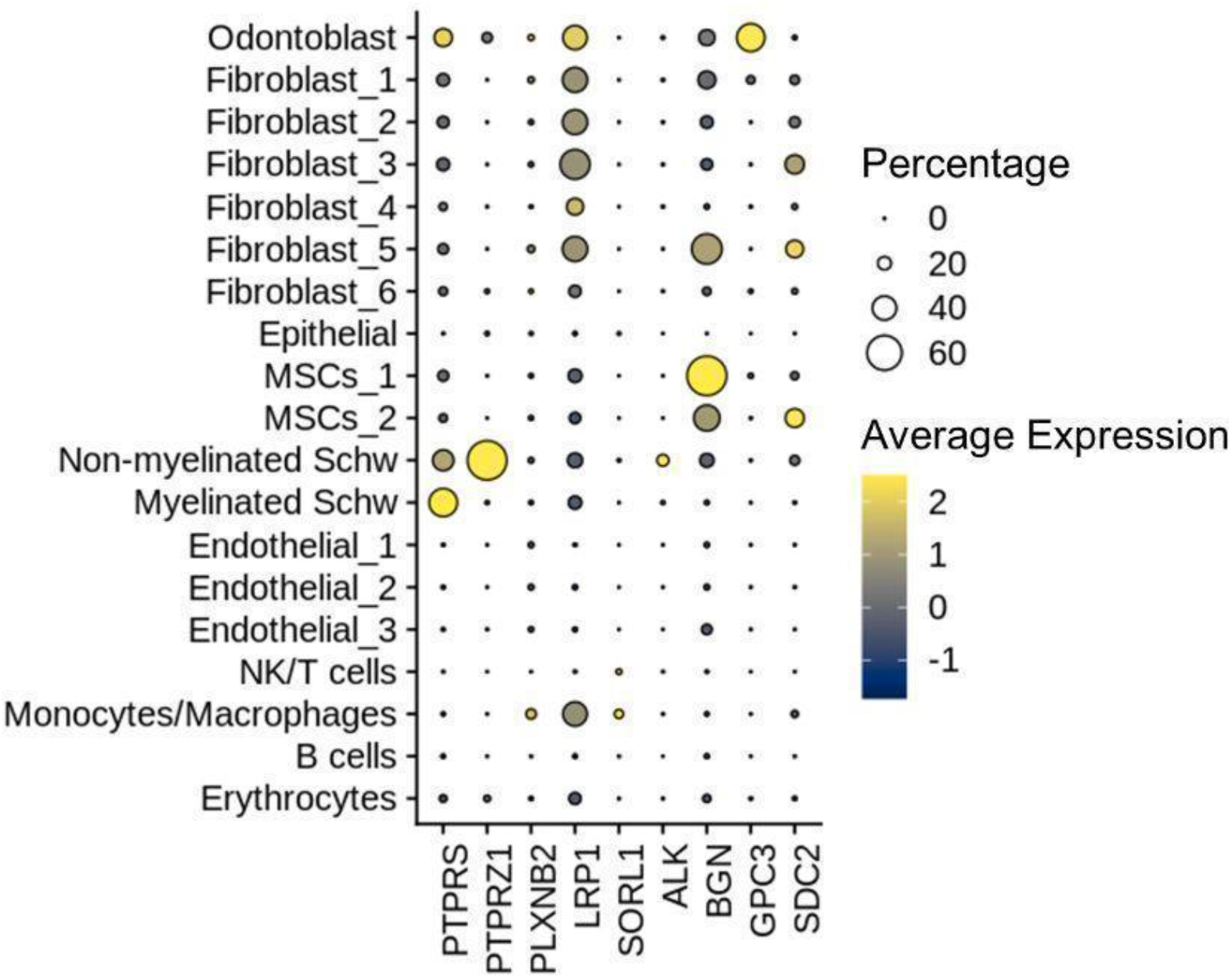
Percentage and average expression levels of PTN and MDK receptors and genes coded for anchor proteins of heparin.

**Supplementary Table 1.**
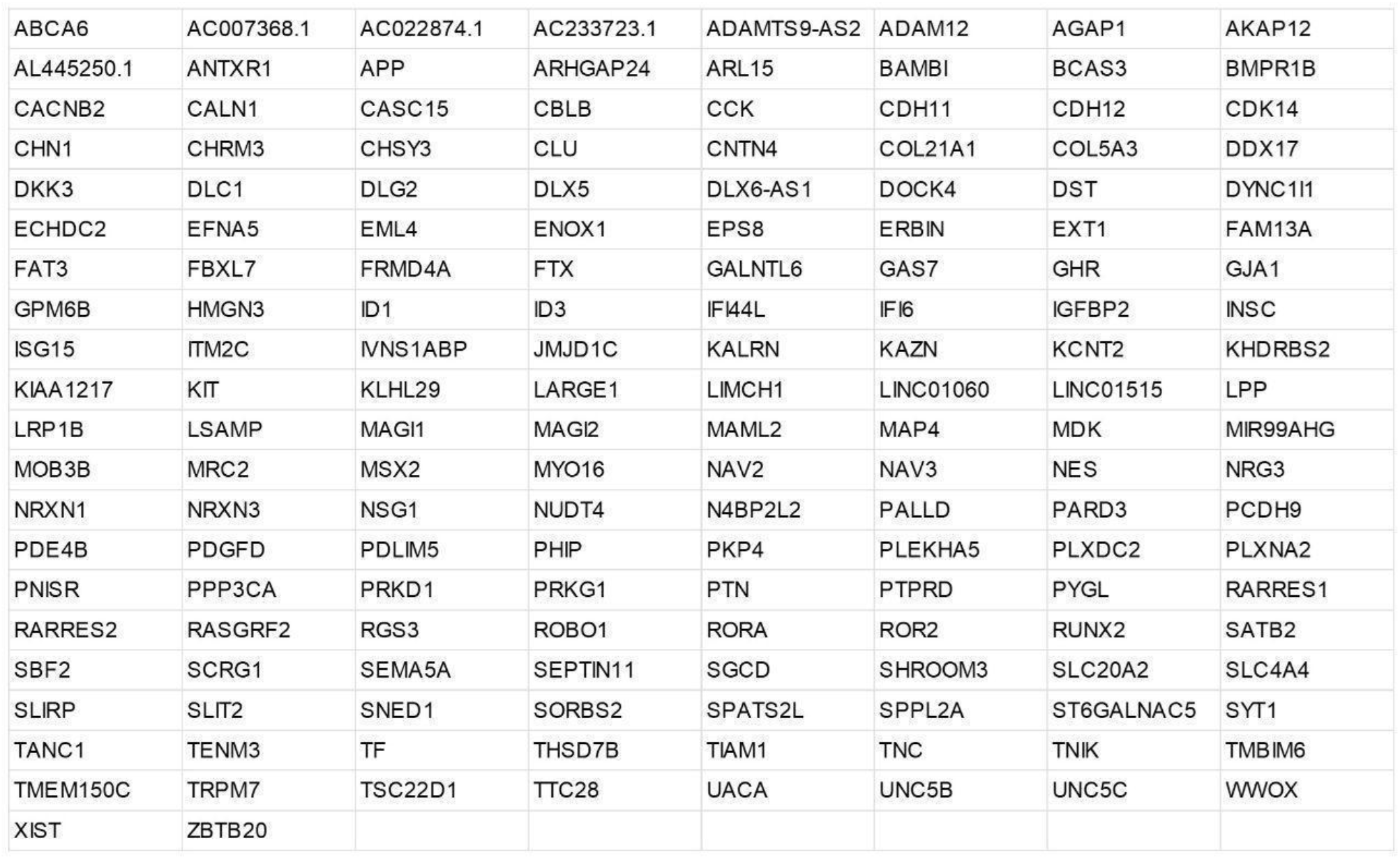

**Supplementary Table 2.**

| **Study** | **Platform** | **Chemistry**<br>**solution** | **Tissue** | **Number of**<br>**datasets** |
| --- | --- | --- | --- | --- |
| GSE185222 | 10x Genomics | 3' V2 | DP | 1 (DP1) |
| GSE161267 | 10x Genomics | 3' V2 | DP | 5 (DP2-6) |
| GSE146123 | 10x Genomics | 3' V3 | DP | 3 (DP7-9) |
| 10X<br>Genomics* | 10x Genomics | 3' V2 | PBMC | 2 |
| PRJEB37166 | 10x Genomics | 3' V2 | BM | 2 |
| GSE155960 | 10x Genomics | 3' V3 | ADP | 2 |
| PRJEB52292 | 10x Genomics | 5' V1 | LUNG | 2 |
| PRJNA754272 | 10x Genomics | 3' V2 | SKIN | 2 |

\* Publicly available datasets from 10X Genomics

